# Active growth signalling promotes cancer cell sensitivity to the CDK7 inhibitor ICEC0942

**DOI:** 10.1101/2021.09.10.459733

**Authors:** G.A. Wilson, G. Sava, K. Vuina, C. Huard, L. Meneguello, J. Coulombe-Huntington, T. Bertomeu, R.J. Maizels, J. Lauring, M. Tyers, S. Ali, C. Bertoli, R.A.M. de Bruin

## Abstract

CDK7 has a central role in promoting cellular proliferation, through the activation of the mitotic CDKs, and by driving global gene expression, through targeting RNA polymerase II. Several recently developed CDK7 inhibitors (CDK7i) have been shown to be non-toxic and to limit tumour growth for a number of cancer cell types and are now in Phase I/II clinical trials. However, the mechanisms underlying the sensitivity of particular cancer cells to CDK7 inhibition remain largely unknown.

To improve the outcome of individual patients and increase the chances of successful CDK7i approval, we assessed which fundamental cellular processes determine sensitivity to CDK7 inhibition, using the highly specific CDK7 inhibitor ICEC0942. Our data shows that selective CDK7 inhibition acutely arrests cells in the G1 phase of the cell cycle, which over time leads to senescence. Through a genome-wide CRISPR knock-out chemogenetic screen we identified active mTOR (mammalian target of rapamycin) signalling, as an important determinant of ICEC0942-induced senescence and show that a cancer-associated mutation that promotes cell growth can increase sensitivity to ICEC0942. Our work indicates that cellular growth is an important predictive marker for sensitivity to CDK7i.

## Introduction

The main driver of cell cycle progression is CDK activity and increased CDK activity has been widely reported in various cancers, making them attractive targets for new treatments (Matthews, Bertoli and de Bruin, 2021). Several studies have shown a marked sensitivity of many cancer types to selective CDK7 inhibition. CDK7 is the CDK activating kinase (CAK), which supports cell cycle progression by phosphorylating and fully activating the main cell cycle CDKs, CDK1, 2, 4 and 6. In addition to its role in CDK activation, CDK7 is also required for RNA polymerase II-dependent transcription through RNA polymerase II (Pol II) and CDK9 phosphorylation (Sava *et al*., 2020).

Cancer cells are sensitive to CDK7 inhibitors at doses at which healthy cells are unaffected, suggesting that these treatments might be well tolerated in a clinical setting. Inactivation of CDK7 results in loss of activation of all mitotic CDKs and is therefore expected to not only prevent cell cycle entry and progression, but also to directly promote cell cycle exit. This suggests that CDK7 inhibitors could trigger cell cycle exit in cancers with either mutations in proteins involved in cell cycle exit (ATM-p53-p16) and/or cell cycle entry pathways (Ras-Raf-Mek-Erk-c-Myc-CyclinD/CyclinE-Rb-E2F), potentially benefiting a wide range of patients. Additionally, with many cancers dependent upon transcriptional drivers, CDK7 inhibitors could hit two birds with one stone: the continued cell cycle progression via CDK inhibition and the addiction to transcriptional drivers via reduced Pol II activity.

ICEC0942 is a recently developed non-covalent, ATP-competitive, selective CDK7 inhibitor. It has been shown to inhibit the proliferation of multiple cancer cell lines at lower concentrations than non-tumorigenic cell lines and to significantly inhibit tumour growth in a mouse colon cancer xenograft model, with no significant side-effects (Patel *et al*., 2018). ICEC0942 is currently in Phase I/II clinical trials for advanced solid malignancies, under the name CT7001 or samuraciclib (NCT03363893) (Sava *et al*., 2019). Importantly, the U.S. Food and Drug Administration (FDA) has granted Fast Track designations to samuraciclib in combination with fulvestrant for CDK4/6i resistant HR+, HER2-advanced breast cancer and samuraciclib in combination with chemotherapy for the treatment of locally advanced or metastatic triple negative breast cancer (Carrick Therapeutics, 2021). With ICEC0942 and other CDK7 inhibitors moving ever closer to the clinic, a more fundamental understanding of what determines sensitivity to CDK7 inhibition is required to optimise the efficacy of these treatments and guide patient stratification.

In this study we investigate how CDK7 inhibition, via ICEC0942 treatment, affects cell cycle entry, progression and exit and what determines CDK7i sensitivity. Our data shows that ICEC0942 limits cell proliferation exclusively by driving cells out of the cell cycle in the G1 phase through induction of senescence, without promoting cell death. Our chemogenetic genome-wide CRISPR screen reveals that active growth signalling, through the mTOR (mammalian target of rapamycin) pathway, promotes ICEC0942-induced senescence. Indeed, we show that mTOR inhibition decreases ICEC0942 sensitivity, suggesting that active growth drives CDK7i-induced senescence. Importantly, we show that reverting a cancer-associated mutation in *PIK3CA*, known to promote cellular growth in breast cancer, to wild-type decreases CDK7i sensitivity. Our data suggests that cancer-associated mutations known to drive cellular growth could be indicative of CDK7i sensitivity and therefore a potential marker to direct their use in the clinic.

## Results

### ICEC0942 inhibits cell proliferation and induces cell cycle arrest in G1

In addition to its role in regulating Pol II-dependent transcription, CDK7 is the CAK, phosphorylating the main cell cycle CDKs, CDK1, 2, 4 and 6. CDK7 inhibition is therefore expected to prevent cell cycle entry and progression and promote cell cycle exit. The selective CDK7 inhibitor ICEC0942 has been shown to be largely cytostatic, inhibiting cancer cell proliferation and tumour growth (Patel *et al*., 2018), but other less selective CDK7 inhibitors, such as THZ1, have been reported to increase cell death (Kwiatkowski *et al*., 2014; Patel *et al*., 2018; Olson *et al*., 2019). To understand how selective CDK7 inhibition by ICEC0942 affects cell cycle progression, we used the non-transformed retinal pigment epithelial cell line RPE1, which retain intact cell cycle check-points. The SRB assay was used to assess cell number upon ICEC0942 treatment and established a GI_50_ of 120nM (Supplementary Fig. S1A). This was confirmed by the MTT viability assay using half, full, and double GI_50_ ICEC0942 concentrations for 48 hours (Supplementary Fig. S1B). At 120 nM of ICEC0942 we observed both a reduction in phosphorylation of the C-terminal domain of RNA po-lymerase II and of the T-loop of CDK1, which are both known to be carried out by CDK7 (Supplementary Fig. S1C).

To investigate if the reduction in cell number upon selective CDK7 inhibition is due to an increase in cell death, as reported for less selective CDK7 inhibitors, such as THZ1 (Kwiatkowski *et al*., 2014; Patel *et al*., 2018; Olson *et al*., 2019), we measured apoptosis by Annexin V/PI staining (Supplementary Fig. S1D). Selective CDK7 inhibition by ICEC0942 only induces a small increase in the percentage of apoptotic cells, which could not account for the strong reduction in cell number observed. This was confirmed by the lack of the apoptotic marker cleaved caspase-3 in cells treated with ICEC0942 (Supplementary Fig. S1E).

With low numbers of apoptotic cells, we next measured if ICEC0942 treatment inhibits cellular proliferation by analysing cell cycle progression via EdU incorporation. EdU is readily incorporated into DNA during S phase, so EdU-positive cells represent the actively replicating cell population. We treated RPE1 cells for 48 hours with ICEC0942 or vehicle, then pulse labelled them with EdU for 1 hour and analysed EdU incorporation by flow cytometry. Treatment with ICEC0942 leads to a striking decrease in the percentage of EdU-positive cells compared to control, with an almost complete loss of S phase cells at higher concentrations (Fig. 1A). Loss of the S phase cell population, upon ICEC0942 treatment, was confirmed by DNA content analysis via propidium iodide staining and flow cytometry. This analysis, which shows the percentage of cells in the G1, S and G2 phases of the cell cycle (Supplementary Fig. S1F) indicates that selective CDK7 inhibition predominantly arrests cells in the G1 phase of the cell cycle.

**Fig 1.**
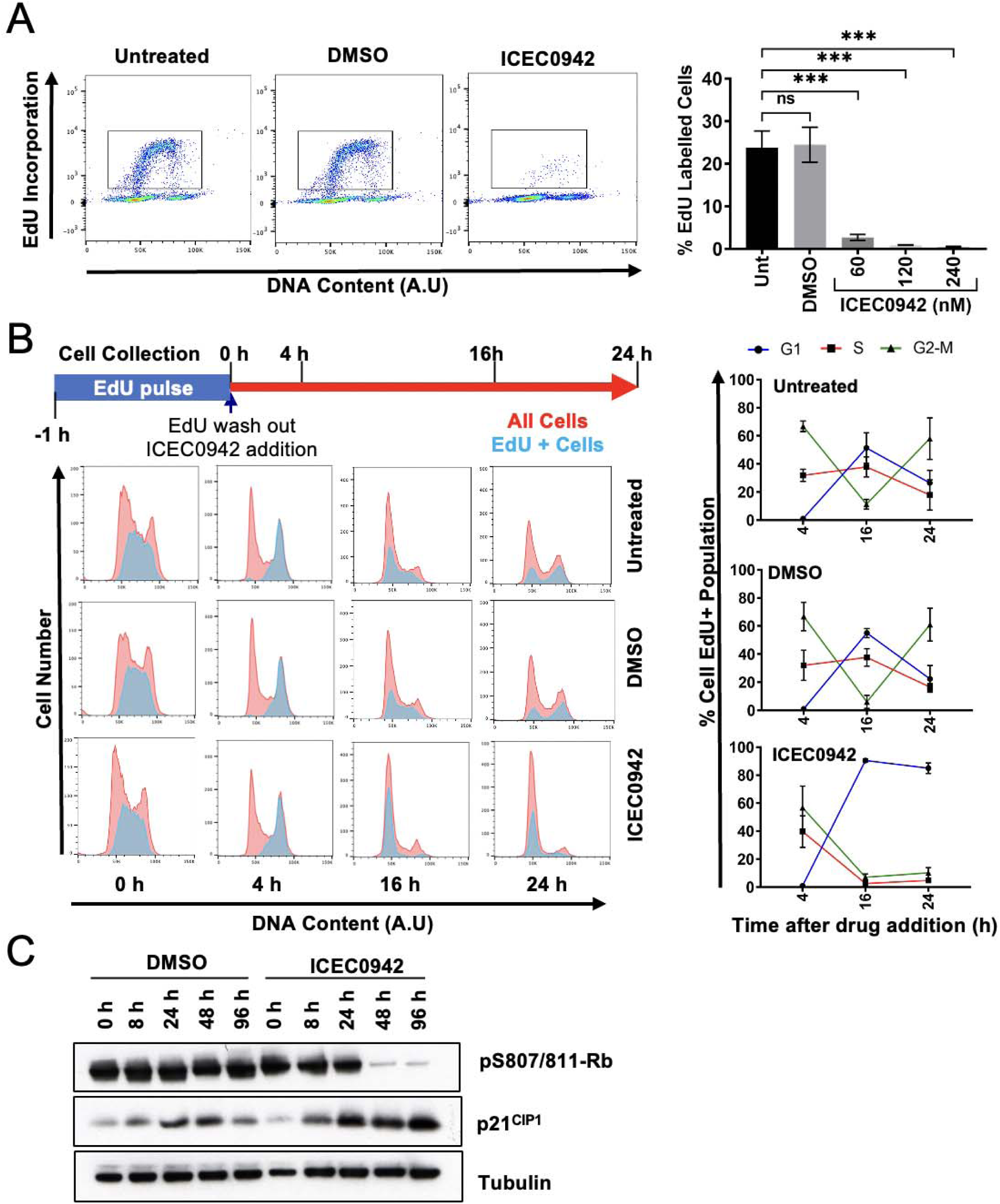
ICEC0942 inhibits proliferation of RPE1 cells by inducing cell cycle arrest in G1. **A**) RPE1 cells were treated with vehicle control or the indicated concentrations of ICEC0942 for 48 hours. Cells were then incubated with EdU for 1 hour before collection. Left: Pseudocoloured plots of flow cytometry data from one representative experiment. DNA content is plotted against EdU incorporation. Inset gate drawn to include EdU+ cells. Right: Quantification of the percentage of EdU+ cells within the population from three independent experiments. Statistical significance is represented by * and was determined using an unpaired two tailed t-test. Error bars show SD. **B)** Top: Schematic illustrating the experimental set up. RPE1 cells were labelled with EdU for 1 hour. Unincorporated EdU was washed out of cells and the cells were treated with vehicle control or 120 nM of ICEC0942. Cells were collected at the indicated time-points after drug addition and then analysed using flow cytometry. Left: Flow cytometry data from one representative experiment. DNA content is plotted against cell count. In red are all of the cells in the sample and overlaid in blue are the EdU+ cells. Right: Quantifications from three independent experiments showing the percentage of EdU+ cells in the different phases of the cell cycle. Error bars show SD. **C)** Western blot analysis from one experiment of whole cell extract collected from RPE1 cells at the indicated time-points after treatment with vehicle control or 120 nM of ICEC0942. Tubulin is the loading control.

As CDK7 is required for the activity of all mitotic CDKs (Lolli and Johnson, 2005; Fisher, 2012), we wanted to establish if ICEC0942 treatment only blocks cells at the G1-to-S transition or also slows down cell cycle progression via a G2 arrest. The S phase population of RPE1 cells were pulse labelled with EdU for 1 hour prior to drug treatment. Unincorporated EdU was washed away before ICEC0942 or vehicle was added and cells were analysed at the indicated time points by flow cytometry (see schematic in Fig. 1B) for up to 24 hours. This allowed analysis of the progression of the S phase cell population, labelled at time-point 0, through the cell cycle during ICEC0942 treatment. In control cells, we observed that the population of cells labelled in S phase complete an entire cell cycle and re-entered S phase at 24 hours. However, whilst ICEC0942-treated cells progressed normally through the G2 and M phases, they collectively arrested with G1 DNA content (Fig. 1B).

These results were confirmed by western blot analysis of protein markers associated with cell cycle progression (Fig. 1C). We observed a decrease in levels of phosphorylated Rb, which coincided with an increase in levels of the CDK inhibitor p21^CIP1^, both of which are well-known markers of cell cycle arrest in G1 (Schwartz and Shah, 2005) (Fig. 1C). This suggests that ICEC0942-treated cells are able to progress through the cell cycle but are unable to enter a new cell cycle. The levels of the CDK4/6 inhibitor p16^INK4a^ do not increase (Supplementary Fig. S1G), which suggests that cell cycle exit pathways (ATM-p53-p16) are not activated.

### ICEC0942 induces a permanent cell cycle exit via senescence

We next investigated whether ICEC0942 induces a cell cycle arrest or an irreversible cell cycle exit. For this purpose we tested if cells are able to proliferate after ICEC0942 treatment. RPE1 cells were treated with ICEC0942 for 2 days after which the drug, which is not covalently bound, was washed out and cells were allowed to grow for an additional 2 or 5 days in culture (see schematic in Fig. 2A). Cell proliferation was assayed by measuring EdU incorporation via flow cytometry (Fig. 2A). Strikingly, the inhibition of cell proliferation observed upon 2 days of ICEC0942 treatment persists for up to 5 days after the wash-out (Fig. 2A). This suggests that the inhibition of cell proliferation induced by ICEC0942 is the result of a permanent cell cycle exit.

**Fig. 2.**
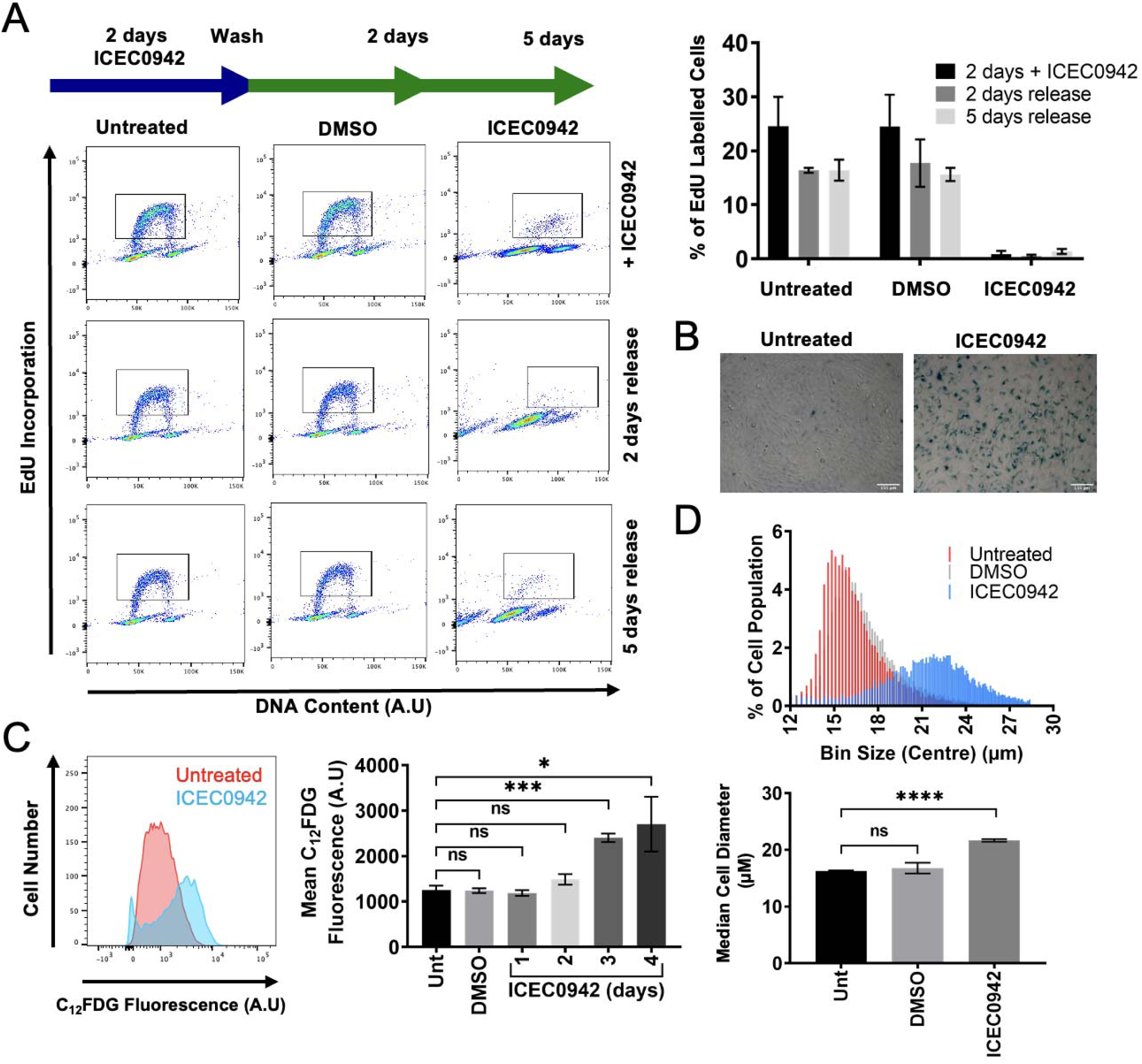
ICEC0942 induces a senescent phenotype in RPE1 cells. **A**) Left: Schematic illustrating the experimental set up. RPE1 cells were treated with vehicle control or 120 nM of ICEC0942 for 48 hours. After 48 hours some cells were labelled with EdU for 1 hour and then harvested. Other cells were washed thoroughly with PBS and given fresh culture media to remove ICEC0942 from the cells. These cells were then allowed to grow in culture for another 2 or 5 days, before undergoing EdU labelling and harvesting. DNA content and EdU incorporation were then measured by flow cytometry. Right: Quantification of the percentage of EdU+ cells within the population from three independent experiments. Error bars show SD. Bottom: Pseudocoloured plots of flow cytometry data from one representative experiment. DNA content is plotted against EdU incorporation. Inset gate drawn to include EdU+ cells.**B)** Representative images of detection of SA β-gal activity in RPE1 cells with the chromogenic β-gal substrate, X-gal in untreated cells and cells treated with vehicle control or 120 nM of ICEC0942 for 4 days. Scale bar represents 155 µm. **C)** Left: Flow cytometry data from one representative experiment, detecting SA β-gal activity with the fluorescent β-gal substrate C_12_FDG in untreated cells and cells treated with 120 nM of ICEC0942 for 4 days. C_12_FDG fluorescence is plotted against cell count. Bottom: Quantification of the mean C_12_FDG fluorescence within populations of cells that were untreated or treated with vehicle control or 120 nM of ICEC0942 for up to 4 days from three independent experiments. Statistical significance is represented by * and was determined using an unpaired two tailed t-test. Error bars show SD. **D)** Top: Coulter Counter data from one representative experiment, with RPE1 cells treated with vehicle control or 120 nM ICEC0942 for 48 hours. Percentage of cell population is plotted against various bins containing cells with different diameters in μm. Bottom: Quantification of the median cell diameter within the population from three independent experiments. Statistical significance is represented by * and was determined using an unpaired two tailed t-test. Error bars show SD.

An irreversible exit from the cell cycle is a hallmark of a senescent state. To further assess if ICEC0942 induces a senescent state we investigated additional markers of senescence. A widely used marker of senescence is an increase in Senescence Associated (SA) β-galactosidase activity (Debacq-Chainiaux *et al*., 2009; Cahu and Sola, 2013). Treatment with ICEC0942 for 4 days results in a clear increase in the number of cells showing a blue staining in their cytoplasm, indicative of SA β-galactosidase activity, compared to control cells (Fig. 2B). A more quantitative assessment of SA β-galactosidase activity, using flow cytometry, indicates a significant increase in SA β-galactosidase activity over time in ICEC0942-treated cells compared to control cells (Fig. 2C). In addition, we observe a striking increase in median cell diameter, measured by Coulter counter, in ICEC0942-treated compared to control cells (Fig. 2D), which is another phenotypic characteristic of senescent cells (Herranz and Gil, 2018). Together these data indicate that selective inhibition of CDK7 induces a senescent phenotype.

### Sensitivity to ICEC0942 depends on active mTOR signalling

To gain insight into the mechanisms via which CDK7 inhibition induces senescence we carried out a genome-wide CRISPR knock-out chemogenetic screen. The screen was performed in NALM-6 cells, a B-cell precursor leukemia cell line, which grow in suspension making them particularly suited for this approach. A pooled custom library of 278,754 different sgRNAs termed extended-knockout (EKO), targeting 19,084 RefSeq genes, 20,852 unique alternative exons, and 3,872 hypothetical genes, was used (Bertomeu *et al*., 2018). We confirmed that ICEC0942 is also able to inhibit cell proliferation and cell cycle progression in this cell line (Supplementary Fig. S2A and S2B). Positive interactors (relative enrichment of sgRNAs) represent genes required for ICEC0942 sensitivity, and the induction of a senescent state, and their inactivation are potential resistance mechanisms. Negative interactors (relative depletion of sgRNAs) represent proteins and processes involved in preventing ICEC0942-induced senescence, whose inactivation would increase ICEC0942 sensitivity and improve the outcome of individual patients. The relative abundance of each sgRNA in ICEC0942-treated cells compared to a control cell population was calculated using a RANKS score and this allowed us to identify genes that may have positive or negative interactions with ICEC0942. We carried out gene ontology (GO) enrichment analysis on the genes that were identified as the top 20 positive and negative interactors with ICEC0942 (Supplementary Fig. S2C). For positive interactors, ‘TOR signalling’ and ‘positive regulation of TOR signalling’ were amongst the most enriched GO terms. Strikingly, for negative interactors, ‘negative regulation of TOR signalling’ was amongst the most enriched GO terms (Supplementary Fig. S2D). The absence of regulators of apoptosis amongst positive interactors further supports our data that ICEC0942 induces a senescent phenotype rather than cell death. Together these results indicate that active mTOR signalling is important for sensitivity to selective CDK7 inhibition.

### ICEC0942-induced senescence depends on mTOR signalling

To test whether active mTOR signalling is important for the induction of ICEC0942-induced senescence, we used Torin1, an ATP-competitive mTOR inhibitor, to inhibit the mTOR signalling pathway (Supplementary Fig. S3A). We first assessed if inhibition of mTOR affects the ability of ICEC0942 to block proliferation. We treated RPE1 cells with either ICEC0942 or Torin1 alone or in combination and assayed cellular proliferation by measuring EdU incorporation (Fig. 3A).

**Fig. 3.**
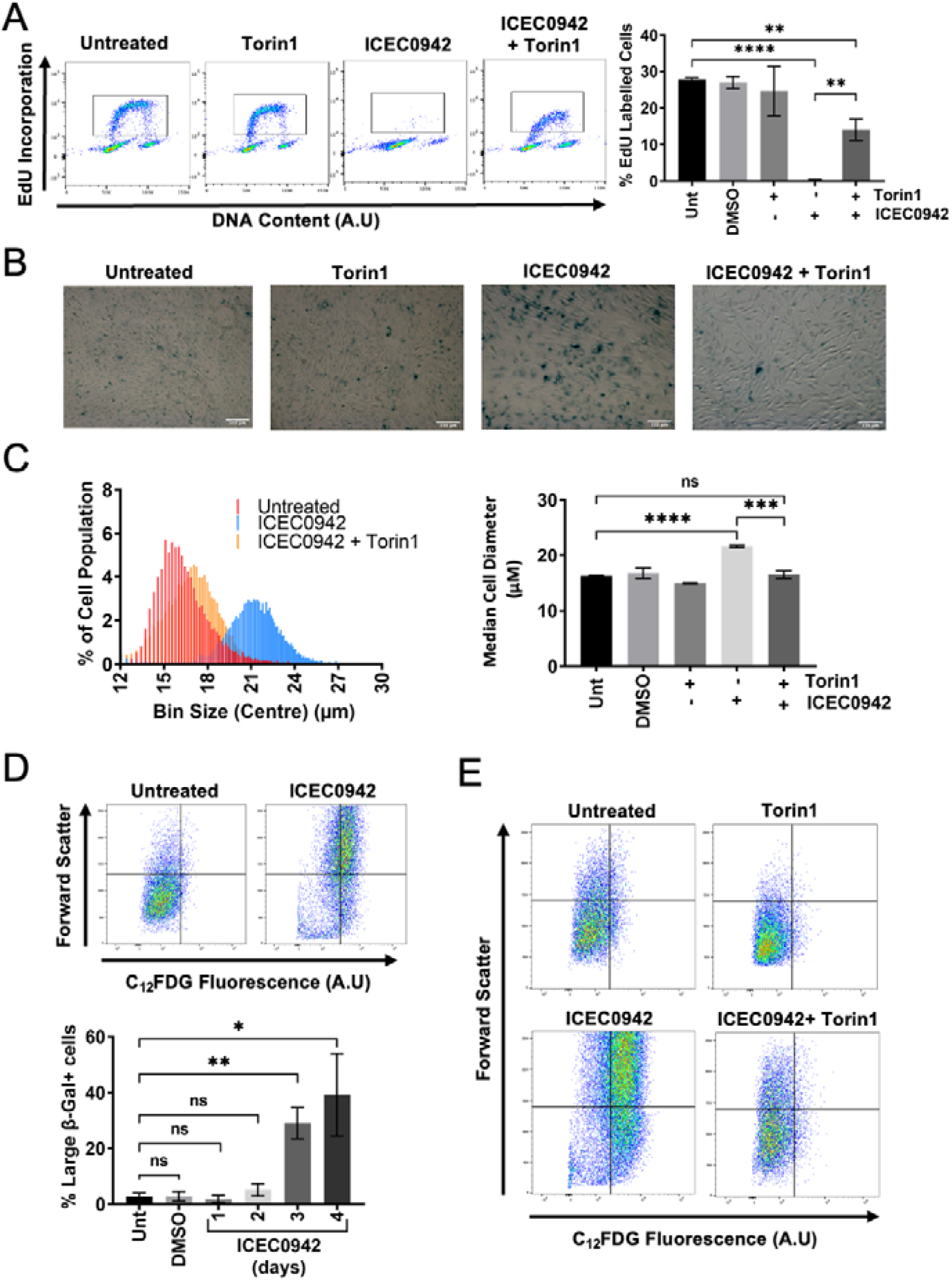
ICEC0942-induced senescence is dependent upon active mTOR signalling in RPE1 cells. **A)** RPE1 cells were treated with vehicle control, 25 nM of Torin1 alone, 120 nM of ICEC0942 alone or 120 nM of ICEC0942 and 25 nM of Torin1 for 48 hours. Cells were then incubated with EdU for 1 hour before collection. Left: Pseudocoloured plots of flow cytometry data from one representative experiment. DNA content is plotted against EdU incorporation. Inset gate drawn to include EdU+ cells. Right: Quantification of the percentage of EdU+ cells within the population from three independent experiments. Statistical significance is represented by * and was determined using an unpaired two tailed t-test. Error bars show SD. **B)** Representative images of detection of SA β-gal activity in RPE1 cells with the chromogenic β-gal substrate, X-gal in untreated cells and cells treated with vehicle control, 25 nM of Torin1 alone, 120 nM of ICEC0942 alone or 120 nM of ICEC0942 and 25 nM of Torin1 for 4 days. Scale bar represents 155 µm. **C)** Left: Coulter Counter data from one representative experiment, with RPE1 cells treated with vehicle control, 25 nM of Torin1 alone, 120 nM of ICEC0942 alone or 120 nM of ICEC0942 and 25 nM of Torin1 for 48 hours. Percentage of cell population is plotted against various bins containing cells with different diameters in μm. Right: Quantification of the median cell diameter within the population from three independent experiments. Statistical significance is represented by * and was determined using an unpaired two tailed t-test. Error bars show SD. **D)** RPE1 cells were treated with vehicle control or 120 nM of ICEC0942 for up to 4 days. Top: Pseudocoloured plots of flow cytometry data from one representative experiment of 4 days of treatment. C_12_FDG fluorescence is plotted against forward scatter, a measure of cell size. Quadrant drawn to distinguish between small, C_12_FDG-cells, small, C_12_FDG+ cells, large, C_12_FDG-cells and large, C_12_FDG+ cells. Bottom: quantification of the percentage of large cells that are C_12_FDG+ from three independent experiments. Statistical significance is represented by * and was determined using an unpaired two tailed t-test. Error bars show SD. **E)** RPE1 cells were treated with vehicle control, 25 nM of Torin1 alone, 120 nM of ICEC0942 alone or 120 nM of ICEC0942 and 25 nM of Torin1 for 4 days. Pseudocoloured plots of flow cytometry data from one experiment. C_12_FDG fluorescence is plotted against forward scatter, a measure of cell size. Quadrant drawn to distinguish between small, C_12_FDG-cells, small, C_12_FDG+ cells, large, C_12_FDG-cells and large, C_12_FDG+ cells.

Combining ICEC0942 and Torin1 significantly increased the percentage of proliferating cells compared to ICEC0942 alone, whilst Torin1 alone showed no effect compared to control. Correspondingly, combined treatment of ICEC0942 and Torin1 led to a decrease in the percentage of cells in G1 compared to ICEC0942 treatment alone (Supplementary Fig. S3B), while increasing the phosphorylation of Rb and loss of p21, indicative of cell cycle progression (Supplementary Fig. S3A). These data suggest that active mTOR signalling is required for the cell cycle block induced by ICEC0942 treatment. To establish if active mTOR signalling is important for the ICEC0942-induced senescent phenotype, we assessed the effect of combined treatment of ICEC0942 and Torin1 on the phenotypic characteristics of senescence previously observed upon ICEC0942 treatment alone. Combining ICEC0942 and Torin1 decreased SA β-galactosidase activity compared to ICEC0942 treatment alone (Fig. 3B and Supplementary Fig. S3C) and also led to a significant decrease in cell diameter, compared to ICEC0942 alone (Fig. 3C). Altogether these data suggest that inhibition of mTOR activity impairs the ability of ICEC0942 to induce a senescent phenotype in RPE1 cells.

### mTOR-dependent cellular growth promotes ICEC0942-induced senescence

The mTOR signalling pathway is known to be a major regulator of cell growth (Thoreen *et al*., 2009). It has been reported that continued cell growth during cell cycle arrest, driven by signalling pathways such as mTOR, can convert a reversible arrest to irreversible senescence (Demidenko and Blagosklonny, 2008; Blagosklonny, 2012; Terzi, Izmirli and Gogebakan, 2016). We tested if mTOR-dependent continued cellular growth is an important factor for ICEC0942-induced senescence. We first assessed if an increase in cell size, as a proxy for cellular growth, upon ICEC0942 treatment correlates with increased SA β-galactosidase activity, a marker of senescence. We measured SA β-galactosidase activity using flow cytometry as in Fig. 2C and plotted it against forward scatter (FSC), which can be used as a measure of cell size (Neurohr *et al*., 2019). We selected large cells based on FSC and compared SA β-galactosidase activity of this population between control and ICEC0942 treatment at various time-points (Fig. 3D, Supplementary Fig. S3D and S3E). We observed that there was an increase in the percentage of RPE1 cells that were large and had increased SA β-galactosidase activity with ICEC0942 treatment compared to control. To test if this increase in cell size and SA β-galactosidase activity required mTOR signalling, we treated the cells with both ICEC0942 and Torin1 (Fig. 3E and Supplementary Fig. S3F and S3G). Inhibition of mTOR signalling combined with ICEC0942 treatment decreases the percentage of large cells with increased SA β-galactosidase activity compared with ICEC0942 alone (Fig. 3E, Supplementary Fig. S3F and S3G), suggesting that increased SA β-galactosidase activity relies in part upon active mTOR signalling.

Altogether these data suggest that an mTOR-dependent increase in cell size correlates with the ability of ICEC0942 to induce senescence more effectively.

### ICEC0942 induces senescence in MCF7 breast cancer cells

Our data shows that ICEC0942 induces a senescent state in non-transformed RPE1 cells, which depends upon mTOR activity. These data suggest that mTOR-dependent cellular growth plays an important role in ICEC0942 sensitivity. Next, we investigated if ICEC0942 also induces a senescent phenotype in cancer cells and if so, whether this is mTOR-dependent. Our previous study showed that ICEC0942 is able to inhibit the proliferation of a number of breast cancer cell lines, including MCF7 cells (Patel *et al*., 2018). We therefore decided to investigate whether ICEC0942 induces senescence in this estrogen receptor-positive breast cancer cell line (Smith *et al*., 2017).

We first tested if ICEC0942 inhibits cell cycle progression in MCF7 cells, as observed for RPE1 cells, via EdU incorporation, as in Fig. 1A. Treatment with ICEC0942 leads to a significant decrease in EdU incorporation in MCF7 cells compared to control (Fig. 4A). Similar to RPE1, we also observed an increase in the percentage of MCF7 cells in the G1 phase of the cell cycle (Supplementary Fig. S4A). Together, this suggests that ICEC0942 can also inhibit cell cycle progression through induction of a G1 cell cycle arrest in MCF7 cells.

**Fig. 4.**
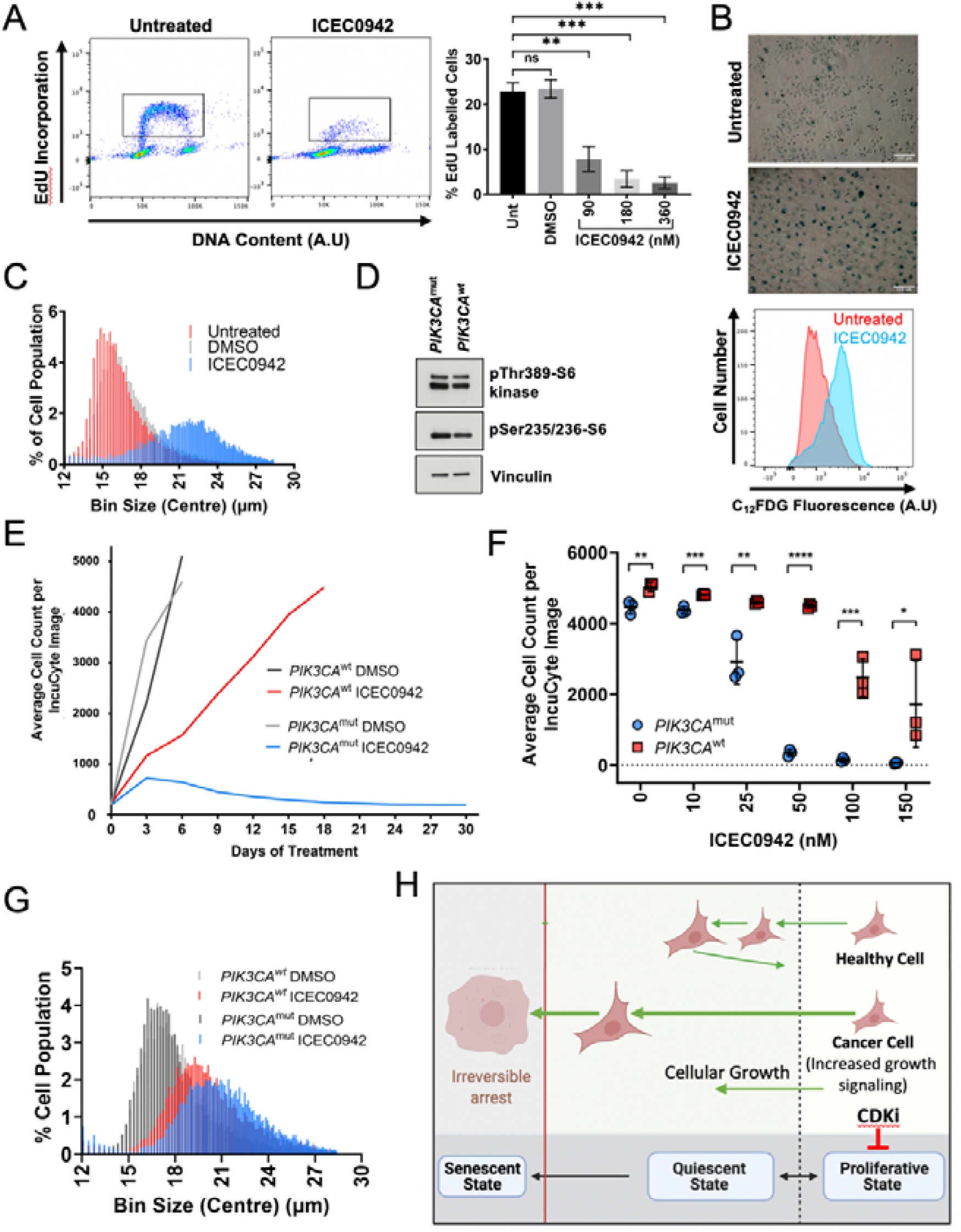
ICEC0942 induces a senescent phenotype more potently in MCF7 cells with activated mTOR signalling. **A**) MCF7 cells were treated with vehicle control or the indicated concentrations of ICEC0942 for 4 days. Cells were then incubated with EdU for 1 hour before collection. Left: Pseudocoloured plots of flow cytometry data from one representative experiment. DNA content is plotted against EdU incorporation. Inset gate drawn to include EdU+ cells. Right: Quantification of the percentage of EdU+ cells within the population from three independent experiments. Statistical significance is represented by * and was determined using an unpaired two tailed t-test. Error bars show SD. **B)** Left: Representative images of detection of SA β-gal activity in MCF7 cells with the chromogenic β-gal substrate, X-gal in untreated cells and cells treated with vehicle control or 180 nM of ICEC0942 for 4 days. Scale bar represents 155 µm. Right: Flow cytometry data from one experiment where C_12_FDG fluorescence is plotted against cell count. MCF7 cells were treated with vehicle control or 180 nM of ICEC0942 for 6 days. **C)** Coulter Counter data from one representative experiment, with MCF7 cells treated with vehicle control or 180 nM ICEC0942 for 4 days. Percentage of cell population is plotted against various bins containing cells with different diameters in μm. **D)** Western blot analysis from one experiment of whole cell extract collected from PIK3CA wild-type and mutant MCF7 cells. Vinculin is the loading control. **E)** PIK3CA wild-type and mutant MCF7 cells were treated with vehicle control or 50 nM of ICEC0942 for 30 days. Cell numbers were assessed by IncuCyte every 3 days. Cell growth is plotted until day 30 or until cells reached confluency. Cell growth curves are from a single representative experiment. **F)** PIK3CA wild-type and mutant MCF7 cells were treated with vehicle control or the indicated concentration of ICEC0942 for 30 days. Cell numbers were assessed by IncuCyte every 3 days. Cell growth is plotted until day 30 or until cells reached confluency. Quantification of average cell numbers after 18 days of treatment from three independent experiments. Statistical significance is represented by * and was determined using an FDR corrected unpaired t-test. **G)** Coulter Counter data from one representative experiment, with PIK3CA wild-type and mutant MCF7 cells treated with vehicle control or 50 nM of ICEC0942 for 4 days. Percentage of cell population is plotted against various bins containing cells with different diameters in μm. **H)** An increase in cell size beyond a particular cell volume can cause uncoupling of protein synthesis with cell size causing a decrease in cytoplasm density, termed cytoplasmic dilution, which is thought to be the basis of irreversible cell cycle arrest. As indicated by our data, active cellular growth (green arrow) during CDK inhibition (red T-junction) might therefore be an important driver of the senescent state, particularly in cancer cells.

Senescence bypass is thought to be an important step in tumorigenesis and a number of genes encoding proteins integral to senescence induction are commonly mutated in cancer (Dimri, 2005). We there-fore tested whether the ICEC0942-induced cell cycle arrest in MCF7 cells is due to senescence, as observed for RPE1 cells. Treatment with ICEC0942 led to a significant increase in SA β-galactosidase activity in MCF7 cells compared to control (Fig. 4B and Supplementary Fig. S4B). Concomitantly, we observed an increase in cell diameter in MCF7 cells treated with ICEC0942, (Fig. 4C), which is in line with our previous observations in non-transformed RPE1 cells. Altogether these data show that inhibition of CDK7 can also induce a senescent phenotype in tumour-derived MCF7 cells.

### A growth-stimulating mutation in MCF7 cells promotes ICEC0942 sensitivity

Our data indicate that cellular growth may be one of the key drivers that determines CDK7i sensitivity. Based on this, cancer-associated mutations that drive cellular growth independently of proliferative signals could be at the basis of the increased sensitivity of cancer cells to CDK7i. MCF7 cells harbour a mutation in *PIK3CA* (E545K), which encodes the catalytic subunit of PI3K. This mutation has been shown to hyperactivate the PI3K/Akt/mTOR pathway, driving cell growth (Liu *et al*., 2018). The *PIK3CA* gene has been shown to be mutated in 20-50% of all breast cancers, and the E545K mutation occurs at one of three “hotspots” frequently mutated in breast cancer (Tian, Li and Zhang, 2019).

To examine the interaction between ICEC0942 and this clinically relevant, growth-promoting mutation, we utilised a pair of isogenic MCF7 cell lines, one with the E545K *PIK3CA* mutation (PIK3CA^mut^) and one where the *PIK3CA* gene has been reverted back to the wild type sequence (PIK3CA^wt^) (Beaver *et al*., 2013). There is increased activation of the PI3K/Akt/mTOR pathway in the PIK3CA^mut^ cells compared to the PIK3CA^wt^ cells, in line with previous data indicating that the E545K mutation induces hyperactivation of this signalling pathway (Fig 4D and Supplementary Fig. S4C).

The PIK3CA^mut^ and PIK3CA^wt^ cells were treated with various concentrations of ICEC0942 for 30 days and cell numbers assessed using an IncuCyte. The number of PIK3CA^mut^ cells was very low throughout the 30 days, indicating that ICEC0942 strongly inhibited the proliferation of these cells (Fig. 4E, 4F and Supplementary Fig. S4D and S4E). ICEC0942 also inhibited the proliferation of PIK3CA^wt^ cells, when compared to control cells, but these cells were less sensitive to ICEC0942 treatment than the PIK3CA^mut^ cells and continued to pro-liferate, particularly at lower concentrations (Fig 4E, 4F and Supplementary Fig. S4E).

The median cell diameter of PIK3CA^mut^ cells was also significantly larger upon ICEC0942 treatment compared to the PIK3CA^wt^ cells at lower drug concentrations (Fig 4G and Supplementary Fig. S4F and S4G), suggesting that increased cellular growth correlates with increased sensitivity to ICEC0942. These data suggest that cells harbouring cancer-associated mutations that promote increased cell growth may be more sensitive to CDK7 inhibition than cells without these mutations.

## Discussion

Here we investigated the mechanism of tumour inhibition and cytostatic effects of the CDK7 inhibitor ICEC0942 to establish the key processes that determine sensitivity to CDK7 inhibition, to guide the use of CDK7 inhibitors and increase the chances of successful CDK7i approval. ICEC0942 was used in our study because of its high selectivity to CDK7 and its non-covalent binding, which allowed the investigation of release and recovery from CDK7 inhibition. Our data indicate that CDK7 inhibition by ICEC0942 induces cell cycle exit through senescence in the G1 phase of the cell cycle, and that this requires active cellular growth. Since growth promoting mutations are present in many cancers, our data suggests that increased cellular growth is likely at the basis of why cancer cells are more sensitive to CDK7 inhibition than normally growing cells.

Strikingly, inhibiting mTOR signalling whilst also inhibiting CDK7 greatly impairs the effect of ICEC0942, limiting cell size by reducing cellular growth, allowing cells to maintain, or return to, a proliferative state. This suggests that anti-proliferation drugs, such as CDK7i but also CDK4/6 inhibitors, should not be used in combination with growth inhibiting anti-cancer drugs as this would prevent a permanent cell cycle exit. It also indicates that, counterintuitively, drugs that promote cellular growth when combined with anti-proliferation drugs could strongly induce permanent cell cycle exit.

Our data shows that larger cells tend to be more senescent than smaller cells, and that reducing cell size, by inhibiting mTOR signalling, decreases the percentage of senescent cells. This supports a model where increasing cell-size beyond a ‘point of no return’ might represent a key driver for inducing a permanent cell-cycle exit (Figure 4H). This is in line with recent work in yeast, which shows that an increase in cell volume beyond a particular point causes ‘cytoplasmic dilution’ and permanent cell-cycle exit (Neurohr *et al*., 2019). More recent work in mammalian cells also indicates that increasing cell size itself results in proteomic changes that gradually approach those found in senescent cells, supporting our hypothesis that cellular growth signalling that drives an increase in cell size *per se* promotes senescence (Lanz *et al*., 2021). Interestingly, this study shows that an increase in cell size promotes senescence in cells where the cell cycle is arrested via the CDK4/6 inhibitor palbociclib. This suggests that CDK7i-dependent senescence is more likely linked to the inhibition of CDK7s role in activating the mitotic CDKs, rather than regulation of RNA polymerase II. Together with our work this indicates that increased growth signalling is an important determinant of the increased sensitivity of cancer cells to mitotic CDK inhibitors (Figure 4H), establishing the potentially central role of cellular growth in driving a permanent exit from the cell cycle in both healthy and malignant cells. Further testing this model, also for other mitotic CDK inhibitors such as CDK4/6 inhibitors, which were recently approved for clinical use, will increase the chances of successful CDK7i approval and guide CDK4/6i clinical use.

## Materials and Methods

### Cell culture and drug treatments

Cell lines used were immortalized hTERT human Retinal Pigment Epithelia 1 (ATCC CRL-4000), NALM-6 cells (a gift from Steve Elledge, HMS) and MCF7 cells (ATCC HTB-22). The pair of isogenic MCF7 cell lines, with and without the PIK3CA mutation, were created by somatic gene targeting to correct the native PIK3CA E545K mutation in *PIK3CA* as previously described (Beaver *et al*., 2013).

Retinal Pigment Epithelia 1 cells were cultured in DMEM/F12 media supplemented with 10% fetal bovine serum, 1% penicillin/streptomycin (Gibco) and 3% sodium bicarbonate (Gibco). NALM-6 cells were cultured in RPMI media supplemented with 10% fetal bovine serum and 1% penicillin/streptomycin (Gibco). MCF7 cells were cultured in DMEM media supplemented with 10% fetal bovine serum and 1% penicillin/streptomycin (Gibco). Cells were treated with ICEC0942 at the specified final concentrations, with Etoposide (Sigma Aldrich, E1383) (0.5 μg/ml), with Bafilomycin A1 (Sigma Aldrich, B1793) 1 hour prior to 5-dodecanoylaminofluorescein di-β-D-galactopyranoside (C_12_FDG) addition (100 nM), with Torin1 ((Merck Millipore, 475991) (25 nM or 5 nM). DMSO (Sigma Aldrich, D2438) was the vehi-cle control and the highest concentration used was 0.0012% of total volume.

MCF7 cell lines were made to stably express nuclear EGFP, by lentiviral transduction of pTRIP-SFFV-EGFP-NLS (Addgene, #86677) packaged in HEK293T cells, to allow for accurate quantification of cell number via IncuCyte Zoom.

### Cell proliferation and viability

To determine the GI_50_ concentration of ICEC0942, cell number in response to drug treatment was assessed using the sulforhodamine B (SRB) assay (Skehan *et al*., 1990). Cells were fixed with 40% Trichloroacetic acid (Sigma Aldrich, T0699) and then incubated with SRB sodium salt (Sigma Aldrich, S9012) (0.4% in 1% acetic acid) for 1 hour at room temperature. After air drying, 10 mM Tris base was added to the cells and absorbance was measured on a VersaMax Tunable Microplate Reader (Molecular Devices). To assess cell viability, the MTT assay was carried out using the MTT Cell Growth Assay Kit (Millipore, CT02) following manufacturer’s instructions. Cells were incubated with the MTT (3-(4,5-dimethylthiazol-2-yl)-2,5-diphenyl tetrazolium bromide) solution for 2-4 hours at 37°C, before the precipitate that formed was dissolved using 0.04 N HCl in isopropanol. Absorbance was measured on a VersaMax Tunable Microplate Reader (Molecular Devices). To determine the response of MCF7 cells expressing nuclear EGFP to drug treatment, plates were imaged using an IncuCyte Zoom and nuclear EGFP was used to assess cell numbers.

### Flow cytometry

For analysis of Annexin V/propidium iodide (PI), cells were collected by trypsinisation and Annexin V staining was carried out using an Annexin V-iFluor 647 Apoptosis Detection Kit (Abcam, ab219919), following manufacturer’s instructions. Cells were trypsinised and washed with PBS. They were then incubated in Assay buffer containing the Annexin-V iFluor 647 conjugate for 30 minutes at room temperature. After this, PI (Sigma Aldrich, P4864-10ML) was added to each sample at a final concentration of 10 μg/ml and the cells were then analysed.

For analysis of EdU/DAPI, cells were trypsinised and resuspended in PBS and then fixed in 4% Formaldehyde (Sigma Aldrich, 252549) for 15 minutes at room temperature. EdU detection was then performed using Click-iT EdU Alexa Fluor 647 Flow Cytometry Assay Kit (Thermo Fisher, C10424) following manufacturer’s instructions. Prior to EdU detection, cells were counted and diluted to the same number in each sample. Cells were washed in 1 mg/ml BSA in PBS, resuspended in Analysis buffer and incubated at room temperature for 30 minutes. After this, cells were incubated in 0.5 μg/ml DAPI (Sigma Aldrich, D9564) for at least 15 minutes at room temperature and then analysed.

For analysis of DNA content by PI staining, cells were trypsinised and fixed in 70% ethanol at -20°C overnight. After centrifugation, the cell pellet was washed with PBS. Cells were then incubated in 350 μl PBS with 100 μg/ml RNaseA (Sigma, R4875) and 50 μg/ml PI overnight at 4°C. For analysis of SA β-gal activity, flow cytometry was carried out as previously described in (Debacq-Chainiaux *et al*., 2009; Cahu and Sola, 2013). Prior to cell collection, the cell’s media was changed and cells were incubated with Bafilomycin A1 at a final concentration of 100 nM for 1 hour at 37°C. After this, the cells were incubated with C_12_FDG (Invitrogen, D2893) at a final concentration of 33 μM for 2 hours at 37°C. The cells were then trypsinised and centrifuged at 300 xg for 5 minutes at 4°C. The cell pellets were resuspended in 500 μl of ice-cold PBS and analysed by flow cytometry.

For all experiments using the flow cytometer, samples were measured on a BD LSRII flow cytometer using DIVA software (BD) and analysed using FlowJo software.

### Western blot

Cell extracts were prepared in RIPA buffer (Tris pH 7.5 20 mM, NaCl 150 mM, EDTA 1 mM, EGTA 1 mM, NP-40 1%, NaDoc 1%), phosphatase inhibitor cocktail 2 and 3 (Sigma P5726 and P0044) 1:1000, and protease inhibitor cocktail (Sigma P8340) 1:1000. Primary antibodies used anti Caspase 3 (Cell Signaling Technology 9662 rabbit 1:1000), anti Phospho-CDK1 (Thr161) (Cell Signaling Technology 9114 rabbit 1:100), anti Phospho-Histone H2A.X (γH2AX) (Ser139) (Cell Signaling Technology 9718 rabbit 1:250), anti p16^INK4a^ (Abcam ab108349 rabbit 1:2000), anti p21^CIP1^ (Cell signalling Technology 2947 rabbit 1:2000), anti Phos-pho-Rb (Ser807/811) (Cell signalling Technology 8516 rabbit 1:1000), anti Phospho-RNA polymerase II CTD repeat YSPTSPS (Ser5) (Abcam ab5131 rabbit 1:5000), anti Phospho-p70 S6 kinase (Thr389) (Cell signalling Technology 9234 rabbit 1:1000), anti Phospho-S6 ribosomal protein (Ser235/236) (Cell signalling Technology 2211 rabbit 1:1000). Secondary antibodies used were goat anti-mouse IgG HRP conjugate (Thermofisher Scientific PA1-74421 1:4000) and goat anti rabbit IgG HRP conjugate (ThermoFisher Scientific 31460 1:4000). GAPDH (Genetex GTX627408 mouse 1:1000), α-tubulin (Cell signalling Technology 3873 mouse 1:20,000) and vinculin (Abcam ab129002 rabbit 1:20,000) were used as loading controls.

### Detection of senescence-associated β-galactosidase activity

For detection of SA β-gal activity, cells were stained using a Senescence β-galactosidase staining kit (Cell Signaling Technology, 9860) following manufacturer’s instructions. Cells were fixed with 1X fixative solution for 15 minutes at room temperature. After washing with PBS, cells were incubated in 1.5 ml of β-galactosidase staining solution overnight at 37°C in the absence of supplemented CO_2_. All images were acquired with a Leica DM IRB microscope using a 10x objective lens.

### Quantification of cell diameter

For analysis of cell diameter, cells were trypsinised and resuspended in culture media. They were then diluted 1:10 in Coulter Isoton II diluent (Beckman Coulter, 8546719) and cell diameters determined using a particle sizing and counting analyser (Multisizer 4 Coulter counter, Beckman Coulter) with Multisizer 4 software.

### Genome-wide CRISPR knock-out chemogenetic screen

#### sgRNAs pooled library generation

The pooled library of 278, 754 different sgRNAs called EKO (extended-knockout) was inserted within the pLX-sgRNA plasmid (Addgene #50662) and lentivirus was generated in HEK-293T cells as previously described (Bertomeu *et al*., 2018). A doxycycline-inducible Cas9 clonal cell line of NALM-6 (using plasmid pCW-Cas9, Addgene #50661) was generated, infected with the pooled lentivirus libraries and underwent blasticidin selection as previously described (Bertomeu *et al*., 2018).

#### Proliferation assay

To determine the IC_30_ concentration of ICEC0942 in NALM-6 cells, a Beckman Coulter BioMek FX robot was used to mix NALM-6 cells with 11 different concentrations of the drugs (1:3 dilutions) and they were then loaded into 384-well plates, keeping solvent concentration constant (0.1%) across all wells (IRIC’s HTS platform, Montréal, Canada). Plates were kept for 3 days in a high humidity chamber installed in a 37 □C 5% CO2 incubator. 50 μl of CellTiter-Glo reagent (Promega) was then added, plates were shaken for 2 minutes and luminescence read on a BioTek Synergy/Neo microplate reader.

#### Chemogenetic screen

A minimum of 70 million cells (250 cells per sgRNA target) of the NALM-6 doxycycline-inducible Cas9 clone were transduced with EKO and treated with 2 μg/ml doxycycline to induce Cas9 7 days before ICEC0942 was added. Prior to drug addition, a minimum of 70 million cells were collected as samples that had had doxycycline added but had not been treated with drugs. The remaining cells were diluted to 400 000 cells per mL and treated with compounds or 0.1% DMSO, 28 million cells for compounds (100 cells per sgRNA) or 70 million for DMSO. In the case of ICEC0942, cells were treated with 350 nM, the estimated IC_30_. A minimum of 70 million cells were left untreated as control. Cell concentration was monitored every 2 days by diluting 1 ml of the cells 1:10 in Isoton II diluent and using a Beckman Coulter Z2 particle count and size analyzer. The cells were diluted accordingly to prevent cells from arresting and compounds or DMSO were added to maintain initial concentrations. After 8 days, the different populations of cells were pelleted, washed with PBS and frozen.

#### Next Generation Sequencing

Genomic DNA extractions were done using the Gentra Puregene Cell kit (Qiagen). sgRNA sequences were amplified by PCR using 462 μg of genomic DNA (corresponding to 70 million cells), 575 μL 10X reaction buffer, 115 μl 10 mM dNTPs, 23 μl 100 μM primer Outer 1, 23 μl 100 μM primer Outer 2, 115 μl DMSO and 145 units of GenScript Green Taq DNA polymerase in a total volume of 5.75 mL or 185 μg (28 million cells; same PCR recipe in 2.5X smaller volumes) as templates. Whenever DNA quantity requirements were not met, everything was used and a corresponding few more cycles performed. Multiple 100 μl reac-tions were set up in 96-well formats on a BioRad T100 thermal cycler and the steps were as follows: 95 □C 5 minutes, 25 cycles of 35 seconds at 94 □C, 35 seconds at 52 □C and 36 seconds at 72 □C, final step of 10 minutes at 72 □C after the last cycle. Completed 100 μl reaction mixes were combined into one tube and vortexed and 1.5 mL aliquots were concentrated to 100 μl by ethanol precipitation.

A second PCR reaction was performed to add Illumina sequencing adapters and 6 bp indexing primers, using 10 μl of 1:20 dilution of unpurified PCR1 product, 10 μl 5X buffer Kapa, 5 μl 2,5 mM dNTPs, 1 μl of PAGE-purified equimolar premix 100 μM TruSeq Universal Adapter 0, 1 μl of 100 μM PAGE-purified TruSeq Adapter with appropriate index, 1 μl DMSO and 5 units Kapa HiFi HotStart DNA polymerase and volumes brought to 50 μl total volume. The steps of the PCR reaction were as follows: 5 minutes at 95 □C, 5 cycles of 15 seconds at 95 □C, 30 seconds at 50 □C and 30 seconds at 72 □C, 5 cycles of 15 seconds at 95 □C, 30 seconds at 56 □C and 30 seconds at 72 □C, followed by 5 minutes final step at 72 □C after the last cycle. A 238-239 bp amplicon was purified using a 1:1 ratio of Spry beads (AxyPrep Mag FragmentSelect-I, Axygen) and 45 bp was read on an Illumina NextSeq 500 with the 23 first bp read in dark cycles (IRIC’s genomic platform, Montréal, Canada). Sequencing coverage was close to 20 samples per lane.

### Data Analysis

Reads were aligned using Bowtie 2.2.5 (Langmead and Salzberg, 2012: PMID: 22388286) in the forward direction only (--norc option) with otherwise default parameters and total read counts per sgRNA tabulated. To generate a single reference sgRNA distribution, read counts from all time-points of the control screens were summed, including untreated and 0.1% DMSO-treated samples. Gene scores were calculated using a modified version of the RANKS algorithm ((Bertomeu *et al*., 2018), Coulombe-Huntington et al., in preparation).

### Statistics

Statistical significance was analysed using Student’s t-tests, unpaired two-tailed. Key throughout: ns=P > 0.05, *=P < 0.05, **=P < 0.01, ***=P < 0.001, ****=P < 0.0001.

Statistical analysis of Gene Ontology (GO) enrichment was carried out using GOrilla, a web-based GO analysis tool (Eden *et al*., 2007, 2009). The REVIGO web server was used to produce a representative sub-set of GO terms (Supek *et al*., 2011). The FDR q-value is the corrected p-value using the Benjamini and Hochberg method (Benjamini and Hochberg, 1995; Eden *et al*., 2007, 2009).

## Acknowledgements

We are grateful to the de Bruin lab members for critical reading of the paper. This work was supported by core funding to the MRC-UCL University Unit (Ref. MC_EX_G0800785) and funded by R.d.B.’s Cancer Research UK Programme Foundation Award.

---

**Supplementary Fig. S1.**
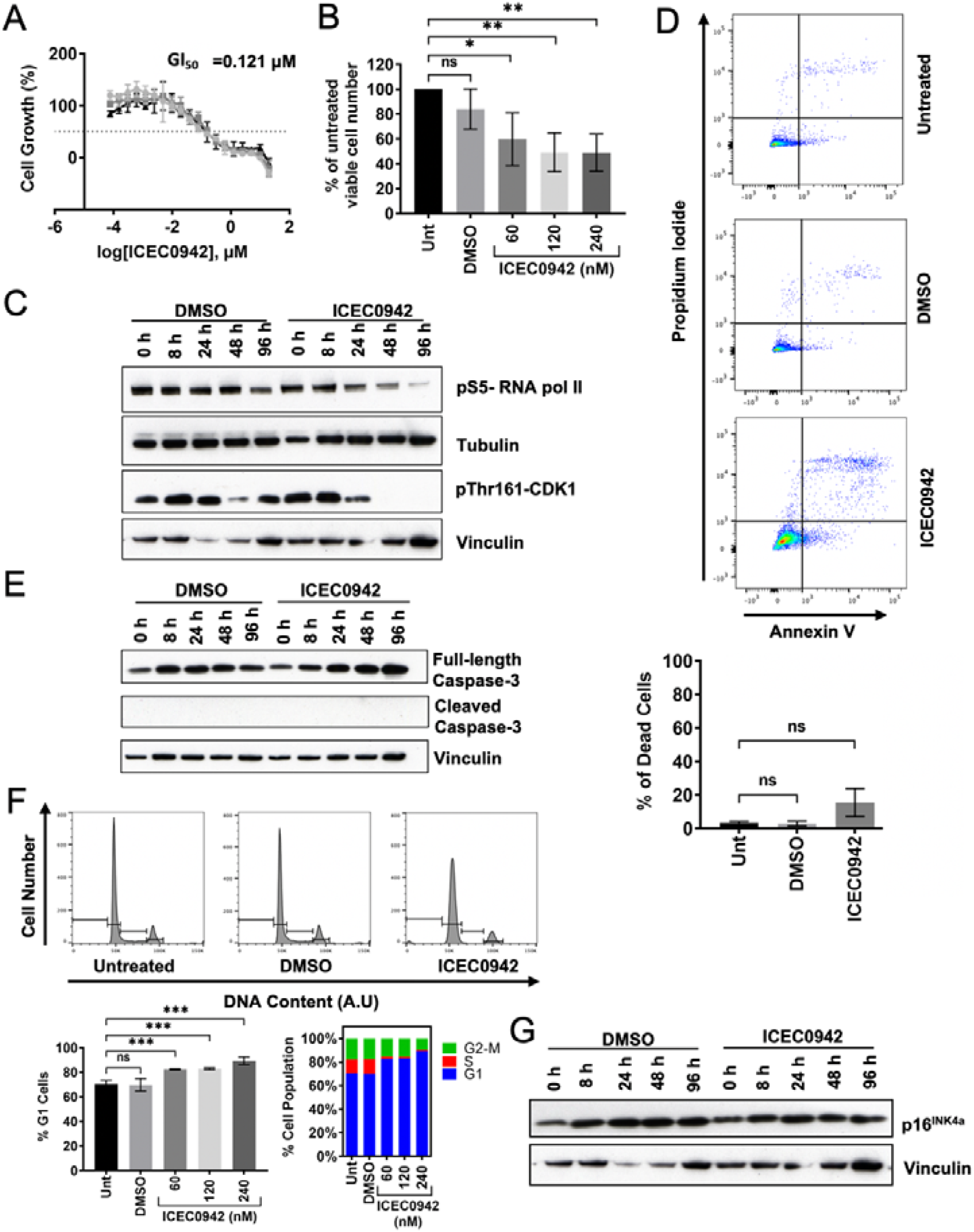
**A**) Dose response curves from three independent experiments measuring the response of RPE1 cells to ICEC0942 treatment using the SRB assay. The GI_50_ is the mean value from these three experiments. Error bars show SD. **B)** Quantifications from three independent experiments showing the percentage of viable cells treated with vehicle control or the indicated concentrations of ICEC0942 for 48 hours compared to untreated cells. Statistical significance is represented by * and was determined using an unpaired two tailed t-test. Error bars show SD. **C)** Western blot analysis from one experiment of whole cell extract collected from RPE1 cells at the indicated time-points after treatment with vehicle control or 120 nM of ICEC0942. Tubulin and vinculin are the loading controls. **D)** RPE1 cells were treated with vehicle control or 120 nM of ICEC0942 for 4 days. Top: Pseudocoloured plots of flow cytometry data from one representative experiment. Annexin V staining is plotted against propidium iodide staining. Quadrant drawn to illustrate the viable cell population (annexin V-, propidium iodide-), early apoptotic cells (annexin V+, propidium iodide-), late apoptotic cells (annexin V+, propidium iodide+) and necrotic cells (annexin V-, propidium iodide+). Bottom: Quantification of the percentage of dead cells within the population (early apoptotic and late apoptotic cells) from three independent experiments. Statistical significance is represented by * and was determined using an unpaired two tailed t-test. Error bars show SD. **E)** Western blot analysis from one experiment of whole cell extract collected from RPE1 cells at the indicated time-points after treatment with vehicle control (DMSO), or 120 nM of ICEC0942. Vinculin is the loading control. **F)** RPE1 cells were treated with vehicle control or the indicated concentrations of ICEC0942 for 48 hours. Top: Flow cytometry data from one representative experiment. DNA content is plotted against cell count. Inset gates were drawn based on DNA content. Left: Quantifications from three independent experiments showing the percentage of viable cells in the G1 phase of the cell cycle. Statistical significance is represented by * and was determined using an unpaired two tailed t-test. Error bars show SD. Right: Quantifications from three independent experiments showing the mean numbers of viable cells in each phase of the cell cycle. **G)** Western blot analysis from one experiment of whole cell extract collected from RPE1 cells at varying time-points after treatment with vehicle control or 120 nM of ICEC0942. Vinculin is the loading control.

**Supplementary Fig. S2.**
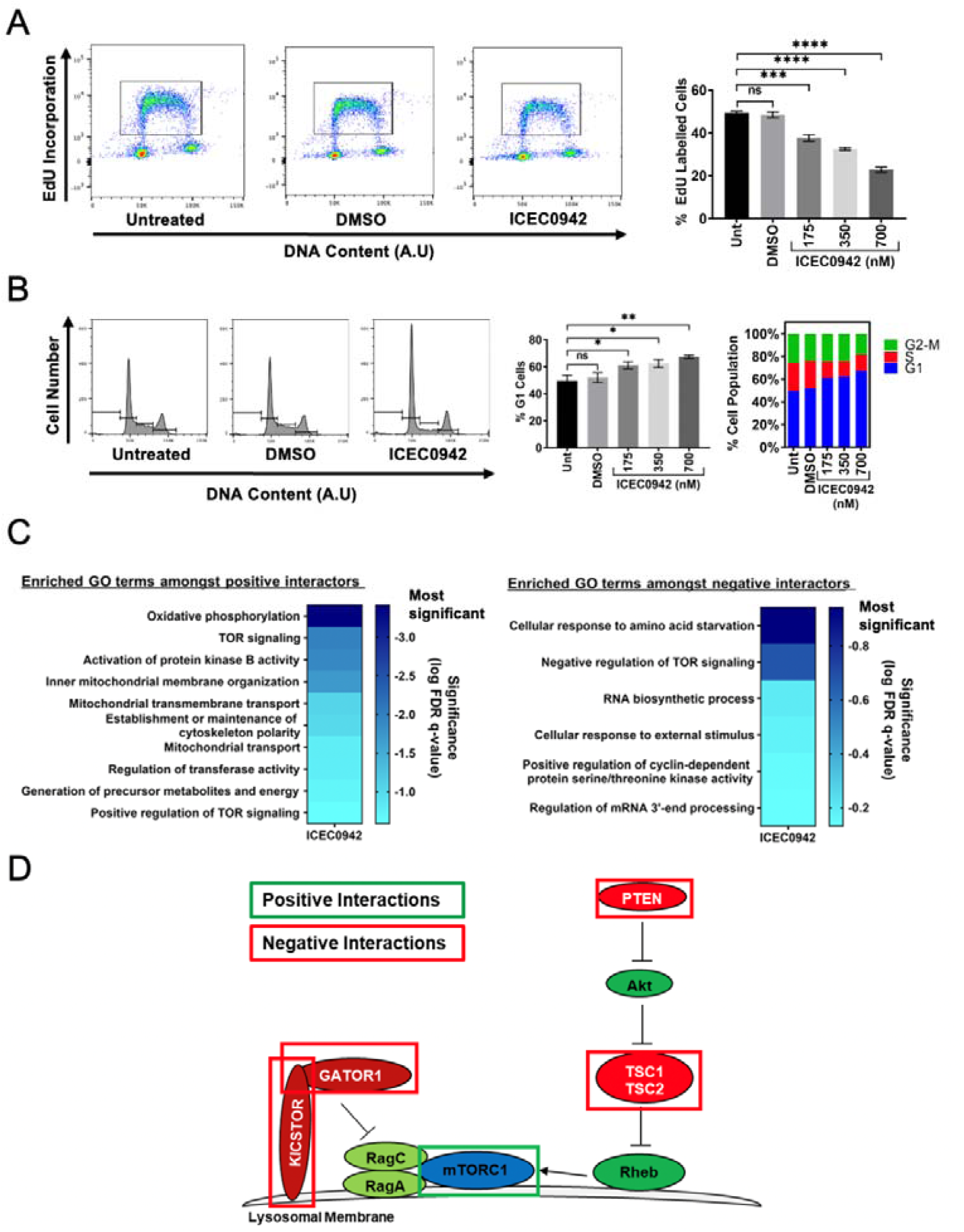
**A**) NALM6 cells were treated with vehicle control or the indicated concentrations of ICEC0942 for 48 hours. Cells were then incubated with EdU for 1 hour before collection. Left: Pseudocoloured plots of flow cytometry data from one representative experiment. DNA content is plotted against EdU incorporation. Inset gate drawn to include EdU+ cells. Right: Quantification of the percentage of EdU+ cells within the population from three independent experiments. Statistical significance is represented by * and was determined using an unpaired two tailed t-test. Error bars show SD. **B)** NALM-6 cells were treated with vehicle control or the indicated concentrations of ICEC0942 for 48 hours. Left: Flow cytometry data from one representative experiment. DNA content is plotted against cell count. Inset gates were drawn based on DNA content. Middle: Quantifications from three independent experiments showing the percentage of viable cells in the G1 phase of the cell cycle. Statistical significance is represented by * and was determined using an unpaired two tailed t-test. Error bars show SD. Right: Quantifications from three independent experiments showing the mean percentage of viable cells in each phase of the cell cycle. **C)** Heat-map representation of the most enriched GO terms amongst the top 20 positive interactors of ICEC0942 (left) and top 20 negative interactors of ICEC0942 (right), as determined using the GOrilla tool (Eden *et al*., 2007, 2009) and processing by REVIGO (Supek *et al*., 2011). These are colour-coded according to the scale on the right, which represents the log (FDR q-value). GO terms closer to the dark blue end of the scale have the most statistically significant FDR q-value and GO terms closer to the light blue end of the scale have the least statistically significant FDR q-value. **D)** Simplified schematic of the mTOR signalling pathway. The boxes indicate components of the pathway that were amongst the top 20 positive interactors of ICEC0942 (green boxes) and the top 20 negative interactors of ICEC0942 (red boxes), as identified by the genome-wide CRISPR knock-out chemogenetic screen.

**Supplementary Fig. S3.**
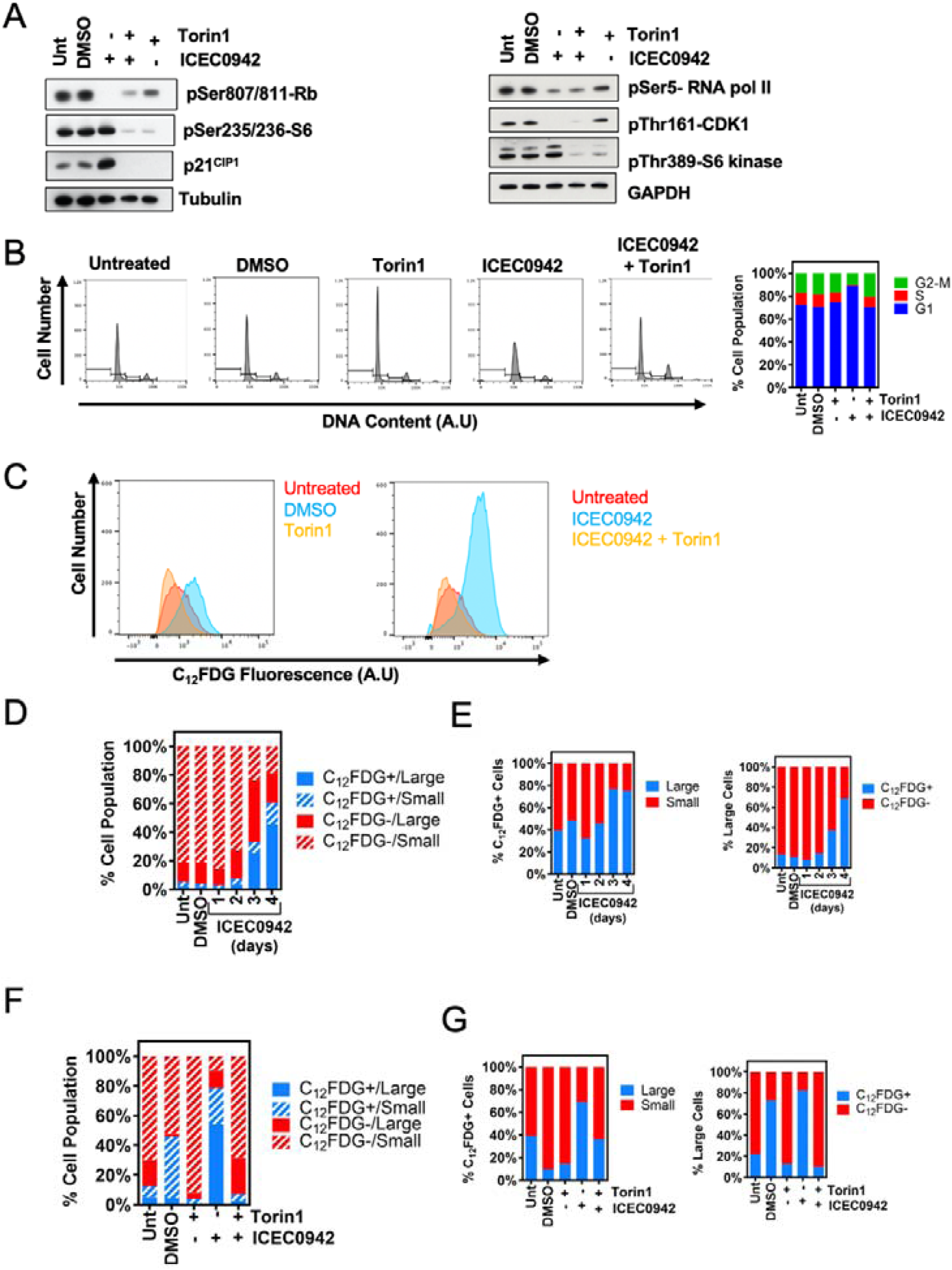
**A**) Western blot analysis of whole cell extract collected from RPE1 cells after 48 hours of treatment with vehicle control, 25 nM of Torin1 alone, 120 nM of ICEC0942 alone or 120 nM of ICEC0942 and 25 nM of Torin1. Tubulin and GAPDH are the loading controls. **B)** RPE1 cells were treated with vehicle control, 25 nM of Torin1 alone, 120 nM of ICEC0942 alone or 120 nM of ICEC0942 and 25 nM of Torin1 for 48 hours. Left: Flow cytometry data from one experiment. DNA content is plotted against cell count. Inset gates were drawn based on DNA content. Right: Quantification from one experiment showing the percentage of viable cells in each phase of the cell cycle. **C)** Flow cytometry data from one experiment, detecting SA β-gal activity with the fluorescent β-gal substrate, C_12_FDG in RPE1 cells treated with vehicle control, 25 nM Torin1 of alone, 120 nM of ICEC0942 alone or 120 nM of ICEC0942 and 25 nM of Torin1 for 4 days. C_12_FDG fluorescence is plotted against cell count. **D)** RPE1 cells were treated with vehicle control or 120 nM of ICEC0942 for up to 4 days. Quantification from one representative experiment showing the percentage of small, C_12_FDG-cells, small, C_12_FDG+ cells, large, C_12_FDG-cells and large, C_12_FDG+ cells. **E)** RPE1 cells were treated with vehicle control, 25 nM of Torin1 alone, 120 nM ICEC0942 alone or 120 nM of ICEC0942 and 25 nM of Torin1 for 4 days. Quantification from one experiment showing the percentages of small, C_12_FDG-cells, small, C_12_FDG+ cells, large, C_12_FDG-cells and large, C_12_FDG+ cells. **F)** RPE1 cells were treated with vehicle control or 120 nM of ICEC0942 for up to 4 days. Left: Quantification from one representative experiment showing the percentages of C_12_FDG+ cells that are large and C_12_FDG+ cells that are small. Right: Quantification from one representative experiment showing the percentage of large cells that C_12_FDG+ and large cells that are C_12_FDG-. **G)** RPE1 cells were treated with vehicle control, 25 nM of Torin1 alone, 120 nM of ICEC0942 alone or 120 nM of ICEC0942 and 25 nM of Torin1 for 4 days. Left: Quantification from one representative experiment showing the percentages of C_12_FDG+ cells that are large and C_12_FDG+ cells that are small. Right: Quantification from one representative experiment showing the percentages of large cells that C_12_FDG+ and large cells that are C_12_FDG-.

**Supplementary Fig. S4.**
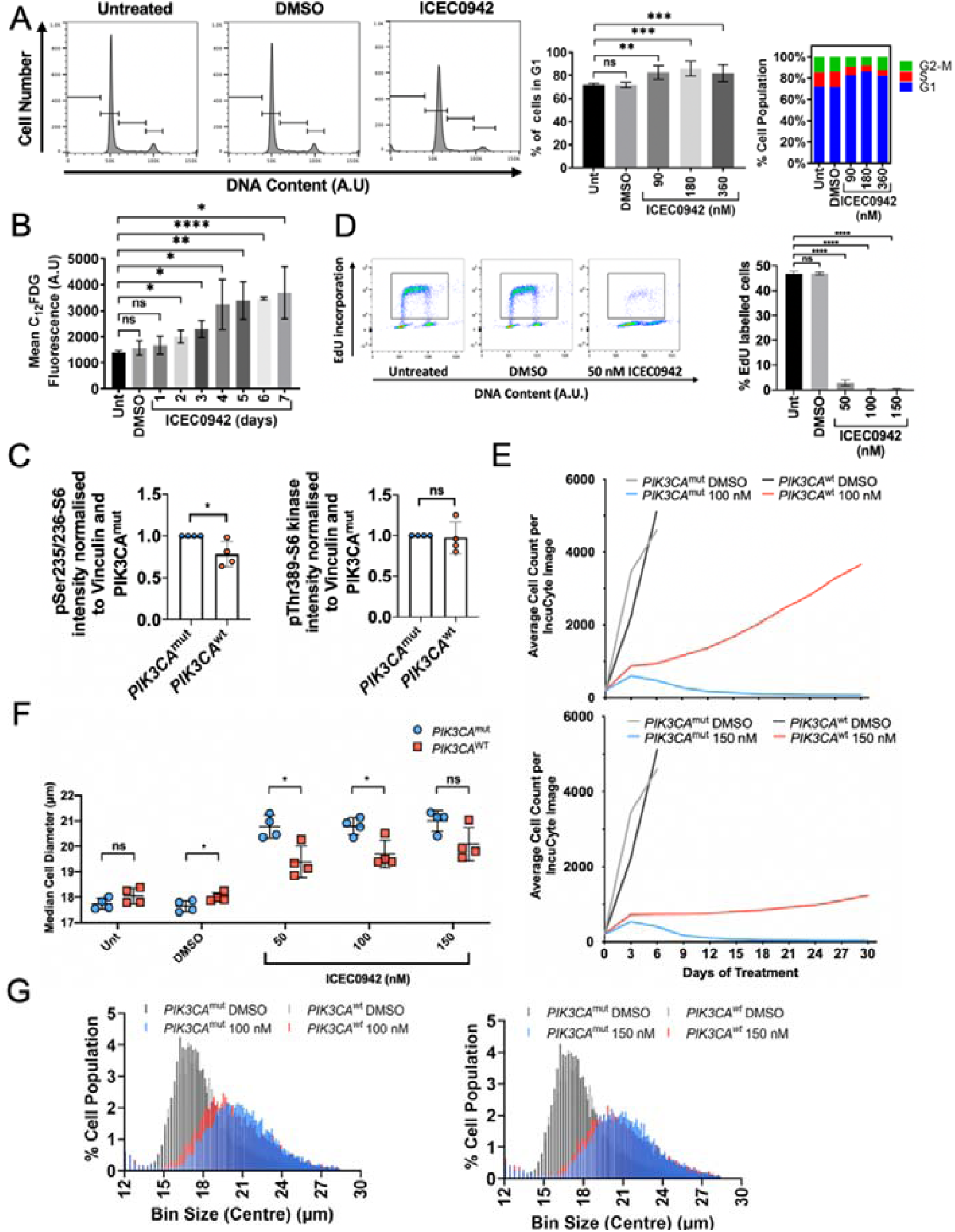
**A)** MCF7 cells were treated with vehicle control or the indicated concentrations of ICEC0942 for 4 days. Left: Flow cytometry data from one representative experiment. DNA content is plotted against cell count. Inset gates were drawn based on DNA content. Middle: Quantifications from three independent experiments showing the percentage of viable cells in the G1 phase of the cell cycle. Statistical significance is represented by * and was determined using an unpaired two tailed t-test. Error bars show SD. Right: Quantifications from three independent experiments showing the mean number of viable cells in each phase of the cell cycle. **B)** Quantification of the mean C_12_FDG fluorescence detected by flow cytometry within populations of cells that were untreated or treated with vehicle control or 180 nM of ICEC0942 for up to 7 days from three independent experiments. Statistical significance is represented by * and was determined using an unpaired two tailed t-test. Error bars show SD. **C)** Quantification of levels of S6 protein phosphorylated at residues Thr389 and Ser235/236, as detected by western blots of whole cell extracts from PIK3CA wild-type and mutant MCF7 cells, from four independent experiments. Statistical significance is represented by * and was determined using an unpaired two tailed t-test. Error bars show SD. **D)** PIK3CA mutant MCF7 cells were treated with vehicle control or the indicated concentrations of ICEC0942 for 4 days. Cells were then incubated with EdU for 1 hour before collection. Left: Pseudocoloured plots of flow cytometry data from one experiment. DNA content is plotted against EdU incorporation. Inset gate drawn to include EdU+ cells. Right: Quantification of the percentage of EdU+ cells within the population. Statistical significance is represented by * and was determined using an unpaired two tailed t-test. Error bars show SD. **E)** PIK3CA wild-type and mutant MCF7 cells were treated with vehicle control or the indicated concentrations of ICEC0942 for 30 days. Cell numbers were assessed by IncuCyte every 3 days. Cell growth is plotted until day 30 or until cells reached confluency. Cell growth curves were from a single representative experiment. **F)** PIK3CA wild-type and mutant MCF7 cells were treated with vehicle control or the indicated concentrations of ICEC0942 for 4 days. Quantification of the median cell diameter within the population from four independent experiments. Statistical significance is represented by * and was determined using an unpaired two tailed t-test. Error bars show SD. **G)** Coulter Counter data from one representative experiment, with PIK3CA wild-type and mutant MCF7 cells treated with vehicle control or the indicated concentrations of ICEC0942 for 4 days. Percentage of cell population is plotted against various bins containing cells with different diameters in μm.

